# Multiple distinct evolutionary mechanisms govern the dynamics of selfish mitochondrial genomes

**DOI:** 10.1101/2022.04.08.487647

**Authors:** Bryan L. Gitschlag, Claudia V. Pereira, James P. Held, David M. McCandlish, Maulik R. Patel

**Author notes:** Corresponding authors* (BLG); (MRP).

## Abstract

Cells possess multiple mitochondrial DNA (mtDNA) copies, which undergo semi-autonomous replication and stochastic inheritance. This enables mutant mtDNA variants to arise and selfishly compete with cooperative (wildtype) mtDNA. Selfish mitochondrial genomes are subject to selection at different levels: they compete against wildtype mtDNA directly within hosts and indirectly through organismal selection. However, determining the relative contributions of selection at different levels has proven challenging. We overcome this challenge by combining mathematical modeling with experiments designed to isolate the levels of selection. Applying this approach to many selfish mitochondrial genotypes in *Caenorhabditis elegans* revealed an unexpected diversity of evolutionary mechanisms. Some mutant genomes persist at high frequency for many generations, despite a host fitness cost, by aggressively outcompeting cooperative genomes within hosts. Conversely, some mutant genomes persist by evading organismal selection. Strikingly, the mutant genomes vary dramatically in their susceptibility to neutral drift. Although different mechanisms can cause high frequency of selfish mtDNA, we show how they give rise to characteristically different distributions of mutant frequency among individuals. Given that heteroplasmic frequency represents a key determinant of phenotypic severity, this work outlines an evolutionary theoretic framework for predicting the distribution of phenotypic consequences among individuals carrying a selfish mitochondrial genome.

## INTRODUCTION

Major evolutionary transitions follow a similar adaptive logic from solitary to social living as from unicellular to multicellular life, prokaryotic to eukaryotic cells, and molecular replicators to cells and genomes.^1-3^ In each case, previously independent entities form a cooperative group that reproduces and undergoes natural selection as a unified collective.^1-5^ By contributing to the fitness of other group members, however, cooperative interactions can select for selfish, “cheater” entities,^6-9^ which occur at each level of biological organization, from the molecular to societal.^9-13^ Although cheating confers an advantage within groups, by assuming the benefits without the costs of cooperation,^6^ cheaters also face disadvantages for numerous reasons. These include suppression or “policing” mechanisms that enforce cooperation, as well as negative frequency-dependent selection, that is, declining fitness as cheaters become more frequent.^12,14^ Selection at higher levels of biological organization, which rely on cooperation, can also counter the fitness advantage of cheaters, consistent with observations ranging from selfish genetic elements to animal societies.^10,14-17^ Thus, different evolutionary mechanisms, such as multilevel and frequency-dependent selection, are thought to help resolve an apparent paradox, namely the coexistence of cooperative and selfish entities.^12,14-17^

Like other major evolutionary transitions, the origin of the eukaryotic cell led to conflicting strategies of cooperation and cheating. This arises from the fact that mitochondria retain their own DNA (mtDNA), a remnant of their bacterial ancestor, whose symbiotic relationship with the nucleus serves as both the foundation of eukaryotic life and as a source of genetic conflicts.^18-20^ In contrast to the typically diploid nuclear genome, an animal cell contains hundreds or thousands of mtDNA molecules,^21,22^ which replicate asynchronously throughout the cell cycle.^23,24^ High mtDNA copy number and relaxed, semi-autonomous replication frequently give rise to a heteroplasmic state, in which a host organism harbors a mixed population of different mtDNA variants. Competition for replication and transmission can select for mutations that confer a selfish advantage over other mtDNA variants within a host, whereas host fitness constitutes a form of group selection favoring cooperative genomes.^18,25-28^ Mitochondria thus serve as an exceptional case study for understanding the dynamics of cooperation and conflict.

Consistent with the conceptual framework of multilevel selection, prior research has uncovered numerous insights into the molecular, physiological, and environmental determinants of the fitness effects of mitochondrial mutations.^29-32^ Many such findings can be credited to the mutant genome *uaDf5* in the model species *Caenorhabditis elegans*.^26,30-39^ Due to a 3.1-kilobase deletion resulting in the loss of four protein-coding and seven tRNA genes (Fig. 1a), *uaDf5* negatively impacts mitochondrial respiration and host fitness.^26,34-36^ Despite these effects, *uaDf5* can persist alongside cooperative (wildtype) mtDNA in a heteroplasmic state for hundreds of generations.^38^ How is this accomplished? In agreement with theory,^40^ experimental work has shown that *uaDf5* propagates not only in spite of but *because* of its adverse effects, by exploiting physiological stress-resistance mechanisms,^32,34,36^ consistent with *uaDf5* being a bona fide cheater.^6^ On one hand, these findings highlight *uaDf5* as a valuable model of a biological cheater. On the other hand, a disproportionate fraction of mechanistic insights have come through studying this genome,^18,26,27,29,30,32,34,36,37,39^ which challenges the ability to draw broad conclusions. Moreover, although prior studies have implicated diverse mito-nuclear interactions in the selfish propagation of mitochondrial mutations,^26,29,32,34,36,41^ as well as substantial variation in their phenotypic effects,^42,43^ this work spans multiple species.^18,27,29^ Because animal species vary widely in ways that affect mitochondrial genome biology, including nuclear genome composition, development and organization of the germline, and environmental context,^29,44^ these numerous confounding factors have strongly limited our ability to draw general conclusions about the evolutionary dynamics of selfish mitochondrial genomes.

**Figure 1:**
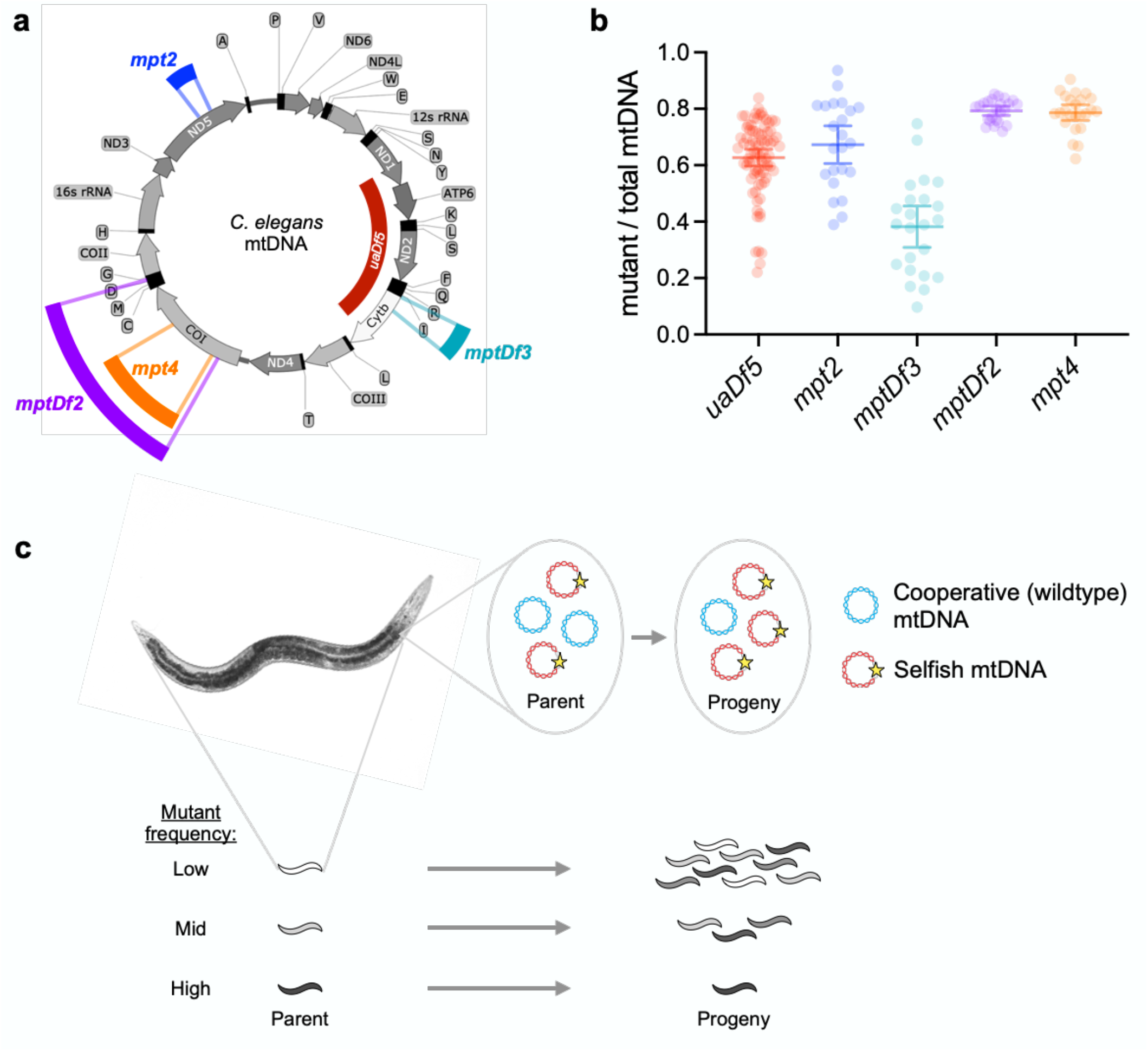
A collection of mutant mitochondrial genotypes stably persisting in a heteroplasmic state despite disruption of essential respiratory genes. **a**, *C. elegans* mitochondrial genome, with the locations and sizes of mutations represented by the color-coded bars. **b**, distribution of heteroplasmic mutant frequencies for age-synchronized adult animals each carrying one of the mutations shown in **a**. Error bars indicate mean and 95% confidence interval of mean. **c**, schematic of multilevel selection, with a selfish mitochondrial genome undergoing positive intra-organismal (within-host) selection and negative organismal (between-host) selection. Shading indicates frequency of selfish mtDNA.

To address these challenges, we developed an approach that leverages a collection of mutant mitochondrial genotypes within a single species, namely *C. elegans*. In particular, we combine mathematical modeling with laboratory experiments to comprehensively measure the evolutionary forces that govern the population dynamics of these stable heteroplasmies. First, we introduce a stochastic population genetic model of selfish mitochondrial genomes, which includes different levels of selection (organismal and intra-organismal) plus genetic drift. Theoretical modeling revealed that different evolutionary mechanisms give rise to qualitatively similar outcomes, such as the persistence of a mutant genome at high heteroplasmic frequencies. Despite these qualitative similarities, however, the distribution of mutant frequencies among hosts contains information about the underlying evolutionary forces. To test our theoretical results, we fit our model to empirical data, obtained from experiments designed to disentangle the levels of selection. We applied this approach to a collection of mitochondrial genotypes, encompassing mutations that affect enzyme complexes I, III, and IV of the electron transport chain (ETC), in addition to the well-known mutant mtDNA variant *uaDf5* (Fig. 1a). Importantly, although each mutant genome stably propagates in heteroplasmic lineages, the distribution of mutant frequencies among heteroplasmic individuals varies considerably between these genotypes (Fig. 1b). Indeed, consistent with our theoretical analysis, we find that although cheating is a common feature of mutant mitochondrial genomes, the evolutionary mechanisms comprising their cheating “strategy,” and which maintain these observed distributions of mutant frequencies, are fundamentally different depending on the affected loci.

## RESULTS

### Population genetic modeling implicates multiple mechanisms in the dynamics of selfish mitochondrial genomes

To understand the complex population genetics of selfish mitochondrial genomes, we developed a population-genetic model that captures natural selection at the organismal and intra-organismal levels, as well as genetic drift within heteroplasmic lineages in order to reflect stochasticity in development and mitochondrial inheritance.

For selfish mtDNA mutants, let us first consider how mutant mtDNA frequency within an individual, *z*, changes between parent and progeny (Fig. 1c, top row). To capture this effect, we define the intra-organismal fitness function *w*_*intra*_(*z*) to be the net relative replication success of selfish versus cooperative (wildtype) mtDNA, per organismal generation, as a function of *z*. Thus, mutant frequency tends to increase when *w*_*intra*_(*z*)>1 but tends to decrease when *w*_*intra*_(*z*)<1. We use a three-parameter fitness function, which can accommodate frequency-dependent or frequency-independent fitness effects and either linear or nonlinear frequency-fitness relationships (see Methods for details). While the intra-organismal fitness function and current mutant mtDNA frequency *z* determine the expected mutant mtDNA frequency among progeny, to capture stochasticity during development and inheritance, we model these dynamics phenomenologically as consisting of a single generation of a Wright-Fisher process where in each generation mtDNA passes through a developmental bottleneck of size *N*. In our modeling framework *N* is best viewed as an effective or “virtual” mtDNA bottleneck size, since it reflects the cumulative effect of drift throughout the organismal lifecycle, both during early development and in the adult germline.

Moving to the organismal level, we treat the inter-organismal dynamics deterministically, corresponding to the case of an infinite population of heteroplasmic hosts. We define organismal fitness *w*_*org*_(*z*) as the expected number of viable offspring per heteroplasmic parent with mutant mtDNA frequency *z*, divided by the expected number of viable offspring per homoplasmic-wildtype parent. We assume a model of purifying selection, so that organismal fitness is monotonically decreasing with increasing selfish mtDNA frequency (Fig. 1c, bottom row), with organisms fixed for the mutant mtDNA becoming sterile or inviable (that is, *w*_*org*_(1)=0), since the mutations investigated here delete one or more essential respiratory gene(s). Because previous heteroplasmy studies have observed threshold effects, whereby modest shifts in mutant frequency can have large phenotypic consequences,^45,46^ we fit a two-parameter fitness function that can accommodate both simple linear and power-law models for how the organismal fitness cost scales with selfish mtDNA frequency, as well as more complicated threshold-like models (see Methods for details).

Combining our organismal and intra-organismal models, each generation begins with a distribution of mutant mtDNA frequencies among heteroplasmic hosts. These frequencies are shifted by intra-organismal selection and the distribution is broadened by intra-organismal drift. Finally, host organisms reproduce with an expected number of viable offspring determined by their mutant mtDNA frequency. As this process repeats across generations, the distribution of mutant mtDNA frequencies among hosts approaches a characteristic shape (given by the dominant eigenvector of a principal submatrix of the transition matrix describing the above evolution process, see Methods), where the shape of this stable mutant frequency distribution reflects the specific parameter values employed. From this distribution, we can calculate the mean fitness of heteroplasmic carriers relative to their wildtype counterparts, as well as the rate of spontaneous reversion to the homoplasmic-wildtype state due to the *de novo* loss of the mutant genome via intra-organismal drift. Together, these values determine the timescale over which an established heteroplasmy will be lost, which can either be short or long depending on the specific parameters. Indeed, under this model we observe that selfish mtDNA can persist in a heteroplasmic state for hundreds, or even thousands, of generations under biologically realistic population genetic conditions.

Examining this model led us to identify a number of distinct evolutionary mechanisms that determine the stable distribution of mutant mtDNA frequencies (Fig. 2). Under the first mechanism, intra-organismal balancing selection, the intra-organismal fitness advantage of the mutant genome vanishes in favor of the wildtype genome at a frequency *z\** that is still too low to encounter significant organismal selection (Fig. 2a,b); that is, there exists a frequency *z\** with *w*_*intra*_(*z\**)=1 and *w*_*org*_(*z\**)≈1. The mutant frequency distribution among hosts is therefore centered at approximately *z*\* and is subject to negligible organismal selection, with genetic drift determining the width of the distribution (Fig. 2c).

**Figure 2:**
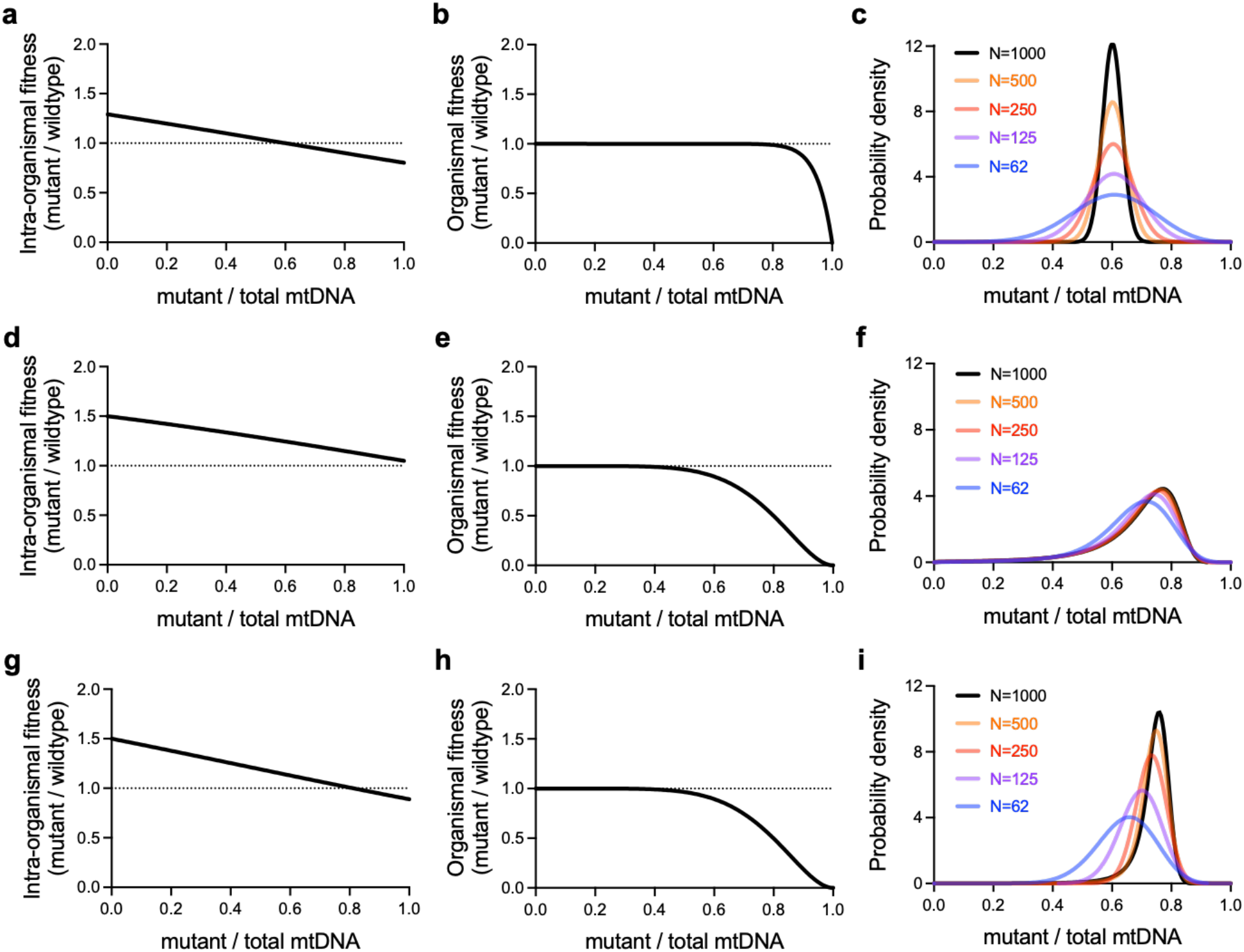
Theoretical modeling reveals the stable maintenance of a selfish mitochondrial genome through a variety of evolutionary mechanisms. **a-c**, the intra-organismal fitness advantage of a selfish mitochondrial genome (**a**) disappears at a frequency too low for the genome to inflict a substantial organismal fitness cost (**b**). The selfish mtDNA is maintained at a mean frequency set by balancing selection at the intra-organismal level, with neutral drift determining the width of the mutant frequency distribution (**c**). **d-f**, a constitutive intra-organismal fitness advantage (**d**) pushes mutant frequency into a range wherein it encounters increasing organismal fitness cost (**e**). Multilevel selection can maintain a broad range of mutant frequencies (**f**), even when neutral drift is minimal. **g-i**, the intra-organismal fitness advantage (**g**) disappears within approximately the same range of mutant frequencies in which organismal fitness declines most rapidly (**h**). Changes in the strength of intra-organismal genetic drift alter both the width and center of the distribution of selfish mtDNA frequencies (**i**).

Under the second mechanism, which could be termed strong multilevel selection, a mutant genome maintains an intra-organismal fitness advantage across all heteroplasmic frequencies, so that there exists no frequency *z\** such that *w*_*intra*_(*z\**)=1 (Fig 2d). Thus, intra-organismal selection constitutively mutant frequency up into a range that elicits a strong organismal fitness cost (Fig. 2e). Under this mechanism, the stable mutant mtDNA frequency distribution is determined predominantly by the forms of the opposing organismal and intra-organismal selection, and is relatively insensitive to changes in the strength of intra-organismal drift, particularly for large mtDNA effective bottlenecks *N* (Fig. 2f).

Finally, in the third mechanism, there exists an intra-organismal equilibrium frequency *z\** with *w*_*intra*_(*z\**)=1, but the organismal fitness cost intensifies rapidly at frequencies similar to *z\** (Fig. 2g,h). Under this mechanism, the shape of the distribution depends on the form of all three forces, namely intra-organismal selection, organismal selection, and drift (Fig. 2i). We therefore call this the mixed mechanism. Together, these findings suggest that mitochondrial mutations can persist at similar heteroplasmic frequencies despite substantial differences in the underlying evolutionary mechanisms. Understanding the mechanisms relevant to any given selfish mitochondrial genome therefore requires that our model be integrated with experiments to measure both selection and drift within hosts, in addition to organismal selection.

### Integrating theory with experiment reveals the evolutionary mechanisms of a selfish mitochondrial genome

As a first application of approach, we inferred the mechanisms determining the form of the stable mtDNA frequency distribution for the well-characterized selfish mitochondrial genome *uaDf5*. To measure intra-organismal selection in the absence of organismal competition, we tracked mutant frequency longitudinally, between isolated individual parents and their respective age-matched progeny (Fig. 3a, see Methods).^26^ At the organismal level, fitness is expected to depend on heteroplasmic frequency, which itself is in flux over the course of organismal development due to intra-organismal selection and drift. This greatly limits our ability to assess the organismal component of selection directly by comparing the fitness of different heteroplasmic hosts. To overcome this challenge and estimate the organismal fitness effects of selfish mtDNA, we instead competed a diverse cohort of heteroplasmic animals against their homoplasmic-wildtype counterparts on the same food plate (Fig. 3b). Since the heteroplasmic cohort is taken from a stock population stably maintaining the selfish mtDNA, the strength of selection against the heteroplasmy reflects the overall fitness of this stable frequency distribution relative to a population lacking the selfish genome, as well as the rate of *de novo* loss of heteroplasmy by genetic drift. Finally, we also collected information on the shape of the stable mutant mtDNA frequency distribution by measuring *uaDf5* frequency in multiple fertile adults sampled from a heteroplasmic stock population (Fig. 3c).

**Figure 3:**
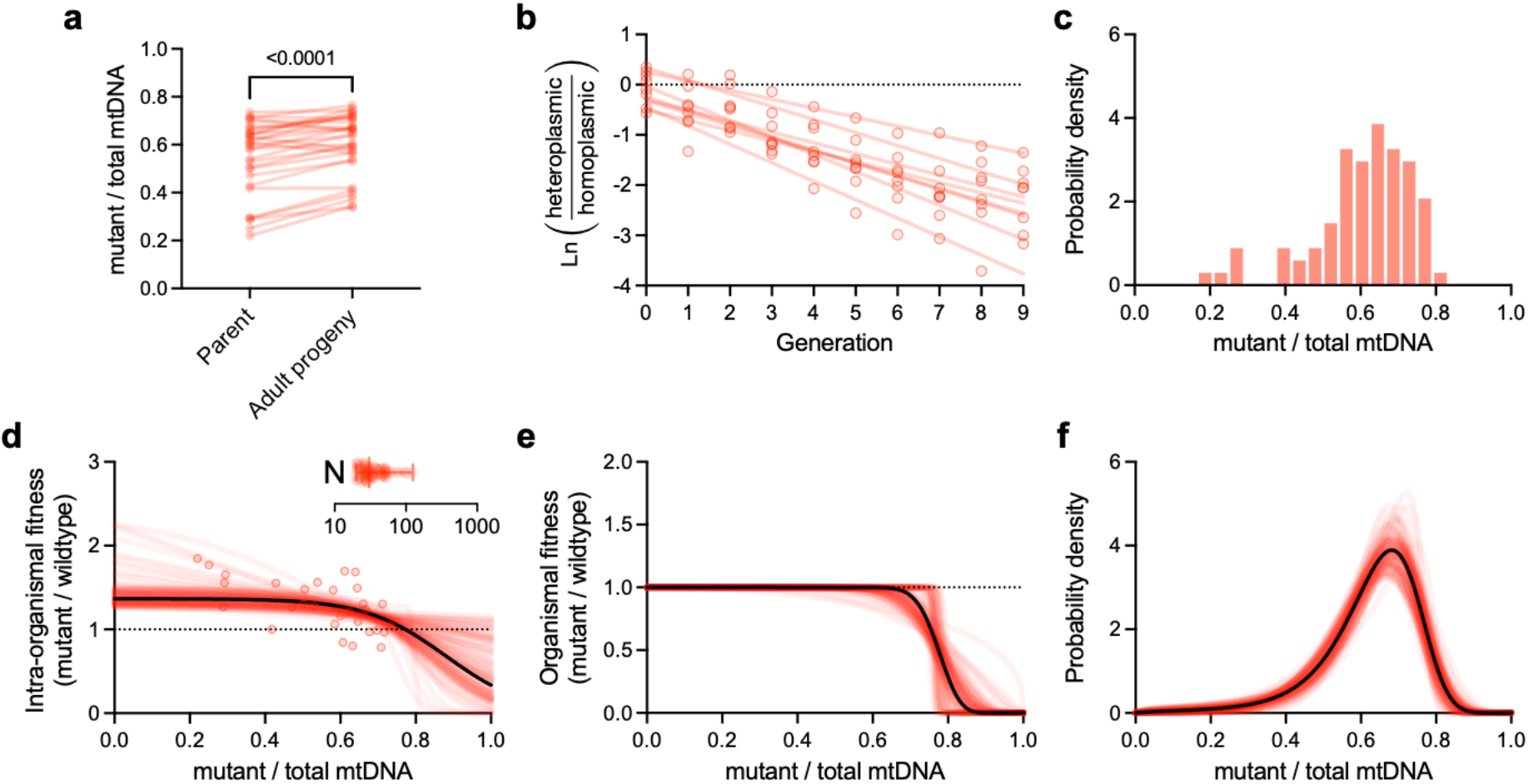
Integration of benchtop experimentation with theoretical modeling reveals the stable maintenance of a selfish mitochondrial genome via a combination of multilevel selection and neutral drift. **a**, pairwise comparisons of mutant frequency (*uaDf5*) between parent and progeny for the purpose of measuring intra-organismal selection on a selfish mitochondrial genome (n=30). **b**, organismal selection against *uaDf5*, measured by directly competing heteroplasmic carriers of the selfish genome against homoplasmic (wildtype) animals on the same food plate, in a mixed population (n=8). **c**, histogram of mutant frequencies (n=81) of age-synchronized adults sampled from a heteroplasmic stock population. Data in **a-c** are taken from our prior study.^26^ **d-f**, maximum-likelihood estimates of the intra-organismal (**d**) and organismal (**e**) fitness effects, each as a function of mutant frequency, and the most evolutionarily stable mutant frequency distribution (**f**). Each plot shows the maximum likelihood estimates corresponding to the empirical data represented in the top row (solid black line) and 100 simulated data sets for parametric bootstrapping (red lines) to visualize the confidence estimates for each plot. Model parameters specifying intra-organismal selection, organismal selection, and intra-organismal genetic drift (**d**, inset, bars indicate mean, lowest and highest values), were collectively estimated using a joint maximum likelihood approach that combines all 3 data sets (top row).

Using the above data (Fig. 3a-c), we inferred the parameters describing organismal fitness, intra-organismal fitness and drift by maximum likelihood, employing a parametric bootstrap procedure to assess uncertainty (Fig. 3d-f, maximum likelihood results in black, bootstrap replicates in red; see Methods for details; Supplemental Fig. 1-3 confirm that our computational procedure can recover the true parameter for simulated data). For *uaDf5*, we found strong evidence for negative frequency-dependent intra-organismal selection (Fig. 3d), where the mutant genome has an intra-organismal advantage when present at low heteroplasmic frequency (*w*_*intra*_(0)=1.37, 95% bootstrap confidence interval 1.27–2.22), and fitness decreases with frequency so that it likely faces an intra-organismal disadvantage at high frequency (*w*_*intra*_(1)=0.34, 95% bootstrap confidence interval 5.8*10^-4^–1.1; *w*_*intra*_(0)>*w*_*intra*_(1) for 100% of bootstraps; *w*_*intra*_(1)<1 for 87% of bootstrap samples). Although our measure of intra-organismal drift is intended to be phenomenological, our maximum likelihood estimate of *N*=29 (Fig. 3d inset, 95% bootstrap confidence interval 20–100) is consistent with previous reports of the estimated effective mtDNA bottleneck in the developing germline.^47^ For organismal selection (Fig. 3d), we infer a threshold-like response, with essentially no fitness cost at low heteroplasmic frequency (*w*_*org*_(*z*)>0.99 for all *z*<0.5 in 92% of bootstrap samples), with the sudden onset of a substantial fitness cost once the mutant reaches sufficiently high frequency (organismal fitness is decreased by half, *w*_*org*_(*z*)=0.5, at *z*=0.77, 95% bootstrap confidence interval 0.75– 0.82). Together, these parameters predict a strongly left-skewed stable mutant mtDNA frequency distribution among hosts (Fig. 3f), as observed in the data (Fig. 3c), a moderate growth defect for the heteroplasmic relative to the wildtype sub-population (relative growth rate of 0.77, 95% bootstrap confidence interval 0.75–0.80), and a low rate of *de novo* loss of heteroplasmy (per-generation probability of mutant loss of 4.1*10^-4^, 95% bootstrap confidence interval 3.7*10^-6^–1.1*10^-3^). Finally, variation in *N* is predicted to substantially impact both the spread and location of the stable *uaDf5* frequency distribution (Supplementary Fig. 4a-c), consistent with the mixed-mechanism hypothesis (Fig. 2g-i). These results suggest that intra-organismal selection, organismal selection, and genetic drift are all important in the dynamics of *uaDf5*.

### Diverse evolutionary mechanisms in the dynamics of selfish mitochondrial genotypes

We sought to compare our results for *uaDf5* with a larger panel of selfish mitochondrial genotypes that exhibit a greater diversity of stable mutant frequency distributions (Fig. 1a,b). These include mutations affecting ETC complex I (*mpt2*), complex III (*mptDf3*), and complex IV (*mptDf2* and *mpt4*), as well as affecting tRNA genes (*mptDf2* and *mptDf3*). We measured selection and collected mutant frequency distribution data (Fig. 4), using the same experimental approach used for *uaDf5*, and applied our theoretical analysis to these data.

**Figure 4:**
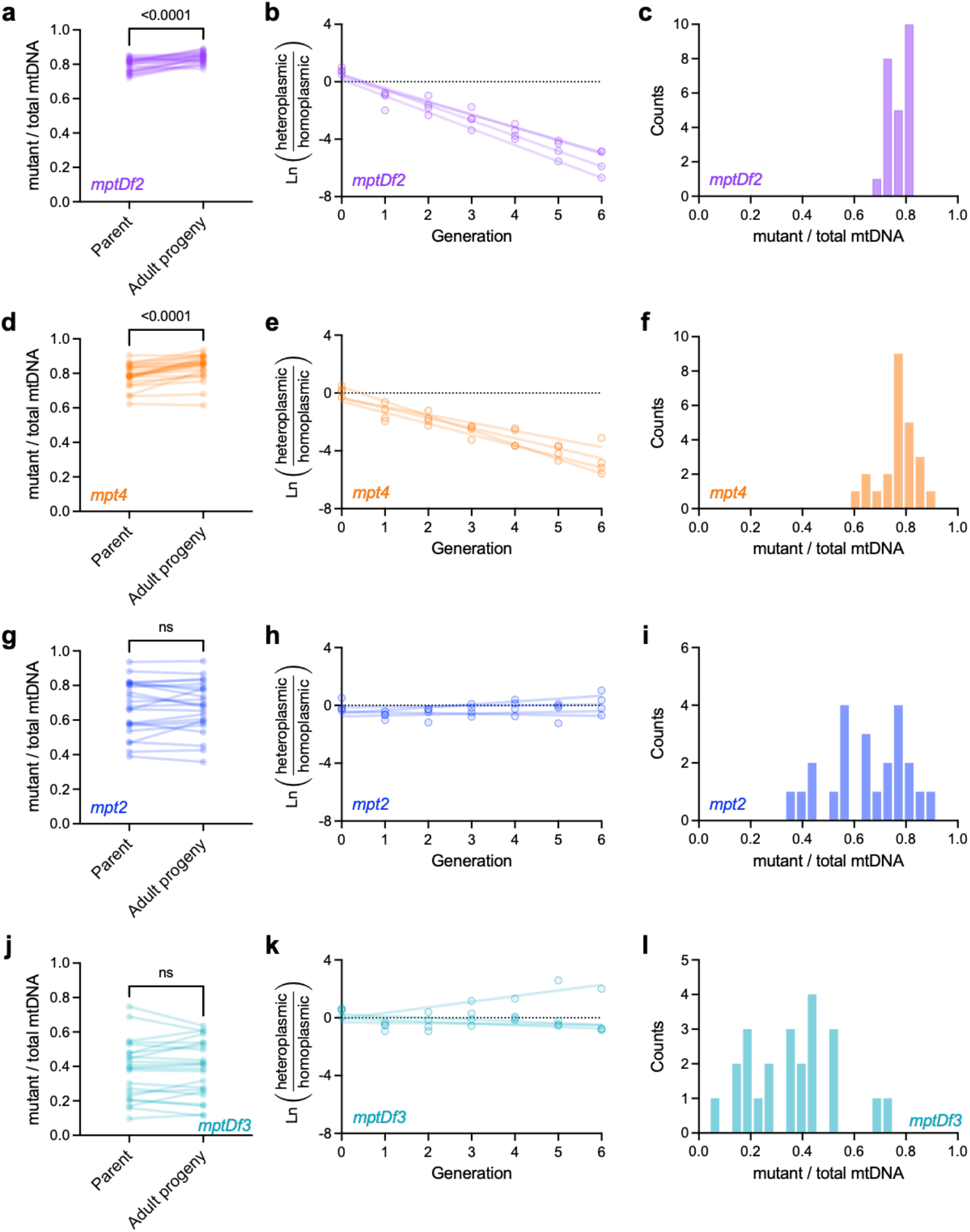
Measurements of intra-organismal selection, organismal selection, and mutant frequency distributions across a panel of putatively selfish mitochondrial genome variants. **a**, *mptDf2* frequencies in isolated parent-progeny lineages (n=24) to measure intra-organismal selection, similar to Fig. 3d. **b**, relative proportions of *mptDf2* heteroplasmic versus homoplasmic-wildtype hosts per generation in populations of animals competing on the same food plate (n=4), to measure organismal selection. **c**, histogram of *mptDf2* frequencies in age-synchronized adults sampled from heteroplasmic stock populations (the same individuals used as parent data points in **a**). **d-f**, similar to **a-c** but for *mpt4* (**d**, n=24). **g-i**, similar to **a-c** but for *mpt2* (**g**, n=23). **j-l**, similar to **a-c** but for *mptDf3* (**j**, n=23).

Overall, we find a surprising diversity of evolutionary mechanisms at play across this panel (Fig. 5), representing all three of the distinct mechanisms described in Fig. 2. For example, although *mptDf2* and *mpt4* have qualitatively similar stable mutant frequency distributions (Fig. 1b), *mptDf2* is governed by the mixed mechanism, similar to *uaDf5*, whereas the distribution for *mpt4* is determined by strong multilevel selection, with little influence of drift. The difference can be seen in Fig. 5a versus Fig 5d, where frequency-dependent intra-organismal selection on *mptDf2* exhibits an internal equilibrium above which selection favors the wildtype (maximum likelihood *z*\*=0.86, 95% bootstrap confidence interval 0.83–0.93) whereas *mpt4* is constitutively favored by intra-organismal selection under the maximum likelihood fit (*w*_*intra*_(*z*)>1 for all *z* for 86% of bootstrap samples, *z\** is greater than any empirically observed *z* in Fig. 4f for 98% of bootstrap samples, and the few bootstrap replicates exhibiting *z\**<1 have *w*_*org*_(*z\**)<0.2). Concordant with this analysis of evolutionary mechanisms, we find that the stable *mptDf2* frequency distribution is sensitive to changes in the mtDNA bottleneck size for (Supplementary Fig. 4f, compare Fig. 2i) whereas the stable *mpt4* frequency distribution is relatively insensitive (Supplementary Fig. 4i, compare Fig. 2f). In contrast, we find that *mpt2* persists primarily through intra-organismal balancing selection (Fig. 5g-i), with a stable mutant frequency distribution centered on *z\** (*z\**=0.68, 95% bootstrap confidence interval 0.62–0.84), where *mpt2* frequency is too low to experience substantial organismal selection (*w*_*org*_(*z*\*)>0.99 for maximum-likelihood estimate and *w*_*org*_(*z\**)>0.92 for 95% of bootstrap samples). Although *mpt2* results are consistent with a negligible role for drift (Fig. 5g and Table 1, 95% bootstrap confidence upper limit of *N*=1,000), we also find evidence of only weak selection at most (Fig. 5h and Table 1, relative growth rate >0.99 for maximum-likelihood estimate and >0.91 for 95% of bootstrap samples). Furthermore, our analysis of the effects of changing the effective mtDNA bottleneck *N* suggests that the width of the stable *mpt2* frequency distribution is highly sensitive to genetic drift (Supplementary Fig. 4l, compare to Fig. 2c). Our modeling results for *mptDf3* are more ambiguous (Fig. 5j-l and Table 1), due to uncertainty in the strength of organismal selection. Like *mpt2* but in contrast to the other genotypes, average *mptDf3* frequency is not significantly different from *z\** (Supplementary Table 7), consistent with intra-organismal balancing selection. Finally, we find no significant organismal selection against *mptDf3* (Fig. 4k and Table 1, relative growth rate 95% bootstrap confidence upper limit >0.99), consistent with the maintenance of *mptDf3* via intra-organismal dynamics. We conclude that although different evolutionary mechanisms maintain the heteroplasmic state, they also give rise to idiosyncratic mutant frequency distributions.

**Table 1:**
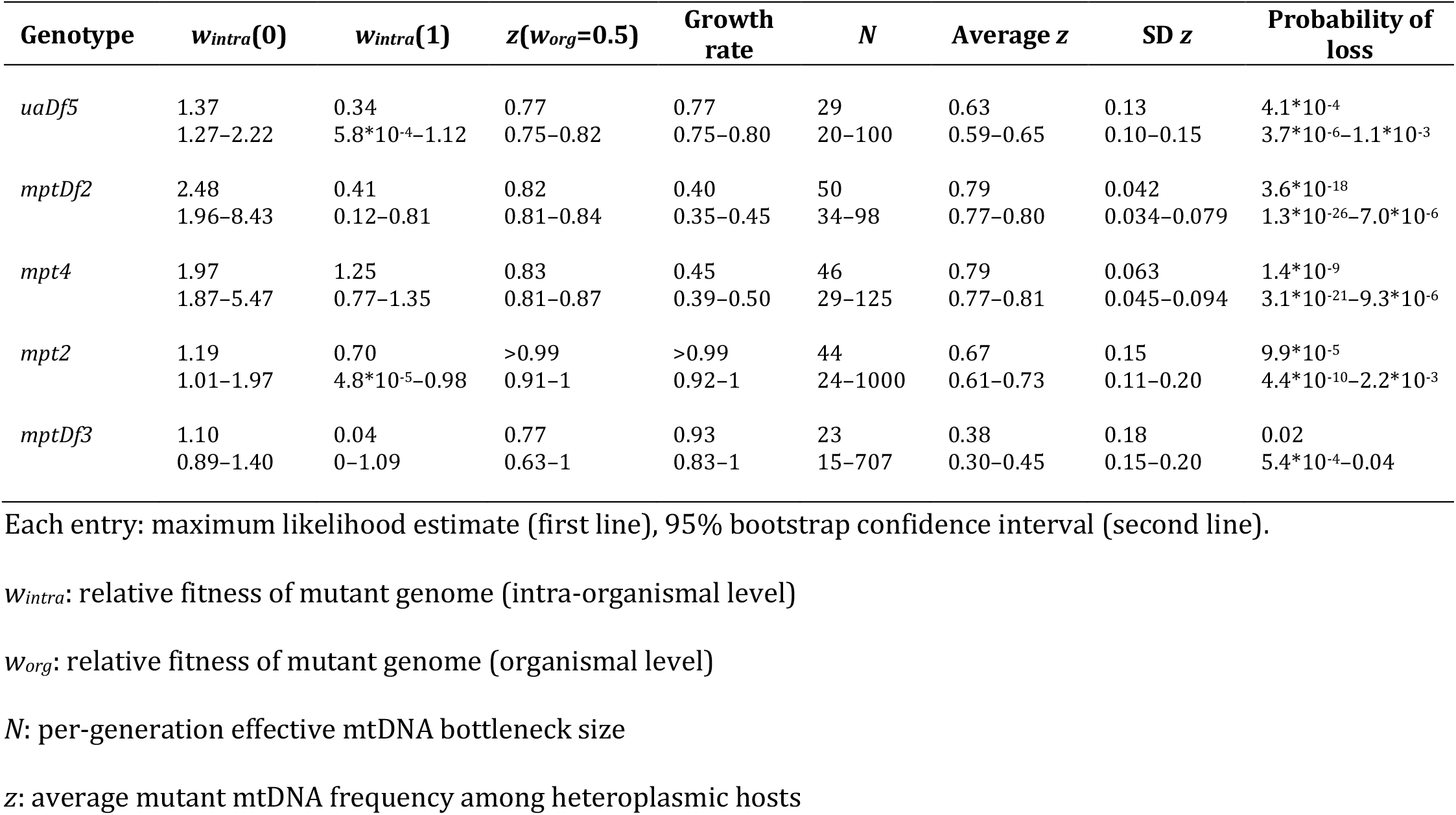
Population genetic characteristics of five selfish mitochondrial genomes.

**Figure 5:**
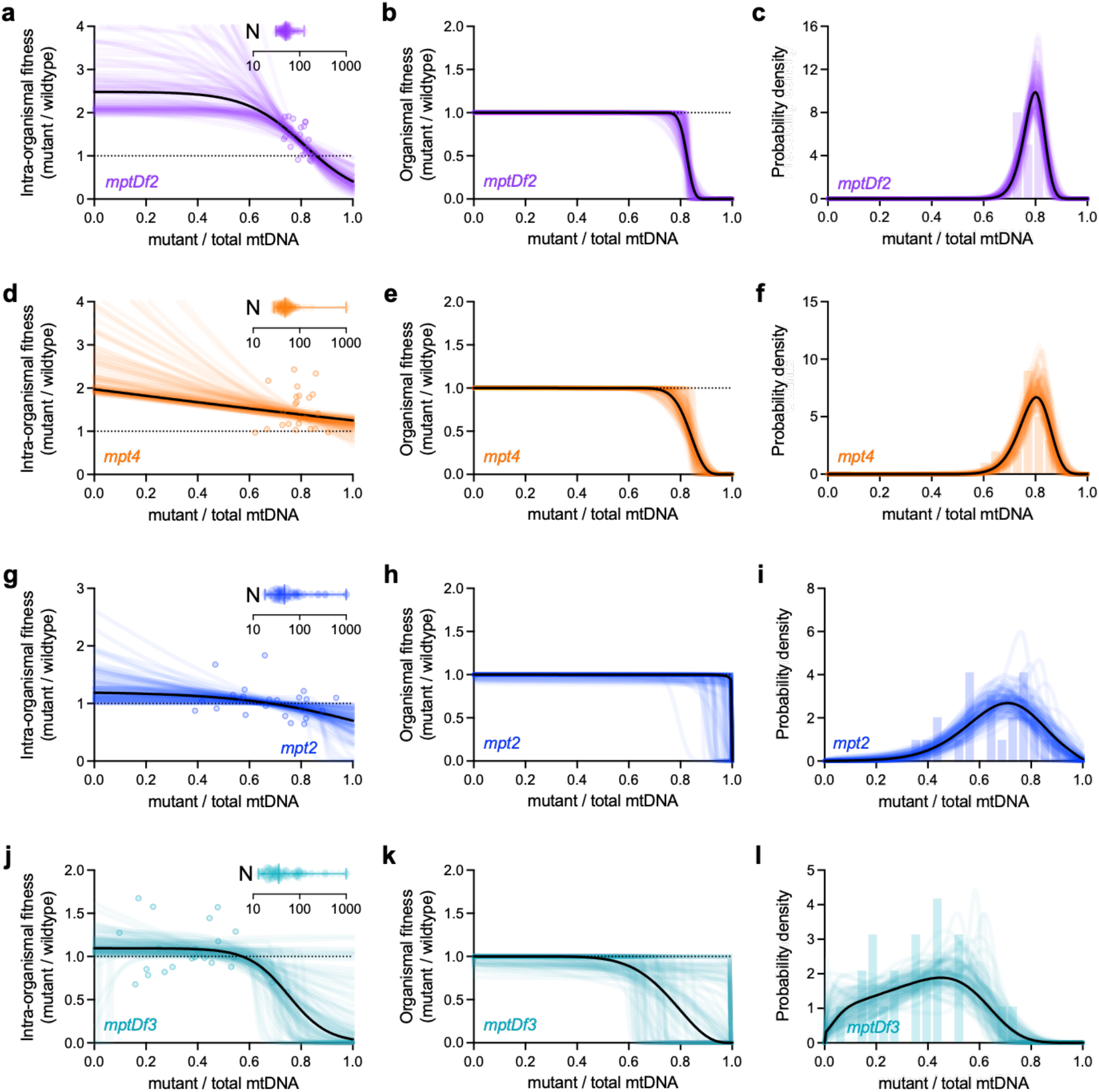
Integrating theoretical modeling with experiment shows the maintenance of selfish mitochondrial genomes in a heteroplasmic state through multiple distinct evolutionary mechanisms. Maximum-likelihood estimates of intra-organismal (left column) and organismal (center column) fitness effects, each as a function of mutant frequency, together with the resulting most evolutionarily stable mutant frequency distribution (right column). Similar to bottom row of Fig. 3, but for the 4 mitochondrial genotypes shown in Fig. 4: *mptDf2* (**a-c**), *mpt4* (**d-f**), *mpt2* (**g-i**), and *mptDf3* (**j-l**). Each plot shows maximum-likelihood estimates corresponding to the empirical data represented in Fig. 4 (solid black line) and 100 parametric bootstrap data sets (colored lines) to visualize confidence estimates. Model parameters specifying intra-organismal selection, organismal selection, and intra-organismal drift (left column, insets, bars indicate mean, lowest and highest values), were collectively estimated using a joint maximum-likelihood approach that combines all 3 corresponding empirical data sets in Fig. 4.

Finally, we sought to explore the potential for long-term maintenance of the heteroplasmic state. We reasoned that high heteroplasmic frequency and a strong intra-organismal advantage (in the case of *uaDf5, mptDf2*, and *mpt4*) limits the probability of *de novo* mutant loss in finite populations, while a minimal organismal cost (in the case of *mpt2* and *mptDf3*) limits the probability of successful invasion by a *de novo* homoplasmic lineage. Using the inferred per-generation probability of mutant loss due to drift (Table 1), together with the organismal fitness benefit of homoplasmic-wildtype state and previously published theory,^48^ we calculated likely persistence times of the heteroplasmic state, defined as the number of generations until the probability of successful invasion is >0.5, for various values of effective organism population size *N*_*e*_. Prior estimates of *N*_*e*_ typically range from 200 to 10^4^ for wild populations of *C. elegans*,^49^ whereas maintenance of laboratory strains involves the frequent transfer of small founding populations to fresh food plates, resulting in an approximate per-generation bottleneck on the order of tens or hundreds of *C. elegans*.^50^ Modeling similar variation in *N*_*e*_, we estimate a wide range of persistence times, reflecting joint uncertainties in organismal fitness and probability of loss (Table 1). Unsurprisingly, our estimates are most consistent with long-term persistence of the heteroplasmic state in small to mid-size populations (Supplementary Table 6). With an *N*_*e*_ of 1,000, for example, our *uaDf5* data are consistent with a median time till invasion on the order of hundreds of generations (95% bootstrap confidence upper limit of 342 generations). Likewise, our results for *mptDf2, mpt4*, and *mpt2* are consistent with a median time till invasion on the order of 10^22^, 10^17^, and 10^9^ generations (95% bootstrap confidence upper limits), respectively. For even smaller populations with an *N*_*e*_ of 100, the data for *mptDf3* are likewise consistent with a median time till invasion on the order of hundreds of generations (95% bootstrap confidence upper limit of 393 generations). We conclude that conditions influencing organism population size on one hand, and conflicting selection pressures on the other, jointly determine which mtDNA mutations are able to persist on long-term evolutionary timescales.

## DISCUSSION

Major evolutionary transitions require cooperative interactions that incentivize group members to assume the benefits of cooperation without the cost of reciprocating.^6-9^ How might cheaters stably persist despite deleteriously affecting the cooperators on which they depend? This question carries broad implications ranging from evolutionary theory to the social and biomedical sciences, as prior exploration of cooperation and cheating spans multiple systems and different levels of scale.^12,14,51^ We sought to systematically compare separate instances of cooperator-cheater dynamics, by leveraging a collection of selfish mitochondrial genotypes existing in a uniform genetic and environmental background.

The selfish proliferation of mitochondrial mutations is responsible for a number of human diseases, affecting an estimated 1 in 4,300 individuals, while an additional 1 in 200 healthy humans carry potentially disease-causing mitochondrial mutations.^46^ The same mutation may thus cause disease in one individual but not another, due to variation in mutant frequency.^45,46^ Consistent with this, disease-causing mutations vary substantially in their ability to propagate from mother to offspring.^43,52^ Predicting the inheritance and development of mtDNA-associated disorders therefore requires a deep understanding of the underlying evolutionary forces, and how they explain observed mutant frequencies. To this end, prior work generally consists of either theoretical modeling or empirical study of mitochondrial mutations,^29,43,53-55^ making it difficult to integrate theory with experiment. Although some modeling studies incorporate empirical data,^29,55-57^ these focus on heteroplasmy dynamics within organisms, or within parent-progeny lineages, making it difficult to combine the levels of selection into a complete evolutionary picture. To address these challenges, we developed a hybrid approach that integrates theoretical modeling with experiments designed to individually probe the levels of selection and the mutant frequency distribution.

The variant *uaDf5* is an exemplary model selfish mitochondrial genome, undergoing positive intra-organismal selection at the expense of host fitness.^26,30,32,34-36,38^ We expand the understanding of *uaDf5* in numerous key ways. Prior studies have found that *uaDf5* proliferates not merely in spite of, but at least partly *because* of, its negative effect on mitochondrial function.^32,34,36^ Conversely, hosts are equipped with mechanisms that limit *uaDf5* proliferation.^32,34,36,37,39^ Together, these findings suggest opposing selection forces at the intra-organismal level, in addition to organismal selection. Consistent with this, and with previous reporting,^26,38^ our modeling results confirm negative frequency-dependent selection for *uaDf5*, with rising frequency leading to loss of its intra-organismal advantage, in addition to an organismal fitness cost. More strikingly, we establish theoretically that the shape of the mutant frequency distribution contains information about the underlying evolutionary forces. Thus, using *uaDf5* data, we show how the combination of multilevel selection and genetic drift explains a previously observed peculiarity, namely the skewed frequency distribution that concentrates most individuals toward the high end.^34^

How generalizable are the results concerning *uaDf5*? To answer this, we expanded our analysis to other stably propagating heteroplasmies, each of which features a mutation known to deleteriously affect at least one essential gene. Like *uaDf5*, some variants persist over time despite a heavy organismal fitness cost, due to a strong intra-organismal advantage. These dynamics can be observed in three of the five heteroplasmies—*uaDf5, mptDf2*, and *mpt4*—and are remarkably consistent with recent mammalian work. For example, some mtDNA mutations in mice and humans rise in frequency from parents to offspring, in a manner consistent with negative frequency-dependent selection,^43^ suggesting a cheating behavior in human mitochondrial disease similar to our findings. For some variants, however, we find no apparent organismal selection, despite deletions in respiratory genes in the mutant genomes. This is true for *mpt2* and possibly *mptDf3*, containing deletions in the genes *ND5* (ETC complex I) and cytochrome b (complex III), respectively, although *mptDf3* results are consistent with a mild organismal fitness cost. How might a lineage lacking an essential gene stably persist with no empirically observed fitness effects? Our modeling shows that a mutant genome evade organismal selection via intra-organismal balancing selection, in which the mutant genome loses its advantage at a frequency where the organismal fitness cost is negligible. One possible molecular basis is that some mutant genomes are more vulnerable to cheater-suppression mechanisms, namely mitochondrial autophagy, consistent with prior empirical work.^34,36,37,39,58-60^ Alternatively, the dynamics of mitochondrial fission and fusion may permit the diffusion of gene products throughout the mitochondrial network, resulting in the genetic complementation of a mutation by nearby wildtype genomes. In either case, such scenarios may enable a mutation to persist in a heteroplasmic state by remaining neutral or nearly neutral at the organismal level.

We note two key limitations of this study, which easily serve as a basis for future work. First, in addition to opposing selection forces, the introduction of selfish genomes by *de novo* mutation, a factor not considered in this study, suggests that selfish mitochondrial genome dynamics represent a persistent phenomenon. Consistent with this, previous mutation-accumulation experiments have identified selfish mtDNA variants in *C. elegans* and the closely related species *C. briggsae*.^25,61^ Together with our study, these findings suggest that a heteroplasmic state may evolutionarily persist by a balance of selection forces, recurring reintroduction of selfish mtDNA, or a combination of these factors.

Conditions beyond the mitochondria represent a second limitation. While our study employs a single host genotype, previous research shows the nuclear genome to influence heteroplasmy dynamics. For example, genetic regulators of mitochondrial biogenesis in response to stress, such as ATFS-1 and FoxO (DAF-16 in *C. elegans*), as well as expression level of mtDNA replication machinery, are important determinants of selfish mtDNA propagation.^26,32,34,36,41^ Moreover, proteins with a more direct role in mitochondrial biogenesis and mtDNA replication, such as POLG and the mtDNA-associated protein TFAM, or that promote binding of replication machinery to mtDNA, reportedly mediate selfish mtDNA proliferation.^30,32,36^ Conversely, genes involved in mitochondrial fission and autophagy reportedly contribute to intra-organismal selection *against* mutant mtDNA.^34,36,37,39,58-60^ Given these findings, we propose that host genotypes and environmental factors, especially those linking physiological stress with mtDNA replication and turnover, will be important determinants of the cooperator-cheater dynamics described here and should be considered in future research.

## METHODS

### Nematode husbandry

*C. elegans* strains used in this study were maintained on 60-mm standard nematode growth medium (NGM) plates (for measuring intra-organismal selection), or 100-mm NGM plates (for measuring organismal selection), seeded with live OP50-strain *E. coli* bacteria as a food source. Nematode strains were incubated at 20°C. In addition to the Bristol wildtype strain and the heteroplasmic *uaDf5* strain featured in prior work,^26^ five additional *C. elegans* strains were used in this study. These consisted of heteroplasmic mutant genomes *mptDf2, mpt4, mpt2*, and *mptDf3*, each crossed into the nuclear background of the Bristol strain.

### Nuclear genome exchange in heteroplasmic strains

To ensure that our analysis was not confounded by variation in the nuclear genome, we completely exchanged the nuclear genome of each heteroplasmic strain with the nuclear genome of wildtype (Bristol strain) *C. elegans*, using a previously published unigametic inheritance method.^62^ Thus, all heteroplasmic strains used in this study have identical nuclear backgrounds. Briefly, each heteroplasmic strain was crossed to the *gpr-1* over-expression strain PD2220 following Mendelian genetics. Next, hermaphrodites of the stable *gpr-1* over-expression heteroplasmy strains were crossed to wildtype males. Non-Mendelian hermaphrodite progeny from these crosses in which the paternal nuclear background is unigametically inherited in the germline cell lineage (determined by pharyngeal mosaic patterning) were individually propagated. Stock strains were established from the progeny of these animals as they have a complete wildtype nuclear genomic background and retain the given heteroplasmy.

### DNA preparation

To prepare animals for quantification of mutant mtDNA frequency, nematodes were transferred to sterile PCR tubes or 96-well PCR plates containing lysis buffer with 100 μg/mL proteinase K. Lysis buffer consisted of 50 mM KCl, 10 mM Tris pH 8.3, 2.5 mM MgCl_2_, 0.45% Tween 20, 0.45% NP-40 (IGEPAL), and 0.01% gelatin, in deionized water. Volume of lysis buffer varied by worm count: 10 μL for individual adults of the parent generation and 20 μL for pooled adult progeny for measuring intra-organismal selection, and 50 μL for pooled animal lysates from the competition experiments for measuring organismal selection. After transferring worms to lysis buffer, each tube or plate was immediately sealed and incubated at -80°C for 10 minutes to rupture nematode cuticles, followed by lysis incubation at 60°C for 60 minutes (90 minutes for pooled nematodes), and then at 95°C for 15 minutes to inactivate the proteinase K. Nematode lysates were then kept at -20°C for stable long-term storage until being used for genotyping and quantification.

### Quantifying mtDNA genotype frequencies

Mutant mtDNA frequencies were quantified as described previously for *uaDf5*,^26^ using droplet digital PCR (ddPCR). Nematodes were lysed as described above and diluted in nuclease-free water with a dilution factor varying depending on nematode concentration: 200x for single adults, 1000x for pooled adults (intra-organismal selection experiment), and 20,000x for pooled nematodes of mixed age (organismal selection experiment). For PCR amplification, 5 μL of diluted lysate was combined with 0.25 μL of 10-μM of each oligonucleotide primer as needed depending on genotype:

*mptDf2*:

Forward 1: 5’-GGATTGGCAGTTTGATTAGAGAG-3’

Reverse 1: 5’-AAGTAACAAACACTAAAACTCCCAAC-3’

Forward 2: 5’-CGTGCTTATTTTTCGGCTGC-3’

Reverse 2: 5’-CTTTAACACCTGTTGGCACTG-3’

*mpt4*:

Forward 1: 5’-CGGTGGTTTTGGTAACTG-3’

Reverse 1: 5’-TCATAGTGTAACACCCGTGAAAATCC-3’

Forward 2: 5’-TGATCCAAGAACTGGAGGTAATC-3’

Reverse 2: 5’-CCTGTTGGCACTGCAATAAC-3’

*mpt2*:

Forward 1: 5’-GAAGAAGGTGGTAGCCTTGAGGAC-3’

Reverse 1: 5’-CGTATAAGAAAAGTCTTGGGATGTTAAG-3’

Forward 2: 5’-GGATTAATTTTCTCAAGGGGTGCTG-3’

Reverse 2: 5’-CTTTTTCAAAGACGAAAACTGTAACC-3’

*mptDf3*:

Forward 1: 5’-CCCTGAAGAGGCTAAGAATATTAGG-3’

Reverse 1: 5’-GGCAATGTCACCAACATCC-3’

Reverse 2: 5’-CCCAATACAATAACTAGAATAGCTCACG-3’

Mixtures of dilute lysate and primer were combined with 12.5 μL of Bio-Rad QX200™ ddPCR EvaGreen Supermix and nuclease-free water to a volume of 25 μL in Eppendorf™ 96-well twin.tec™ PCR plates. Droplet generation and PCR amplification were performed according to manufacturer protocol. Wildtype and mutant-specific primers were combined in the same reaction, and each droplet was scored as containing either wildtype or mutant mtDNA using the 2-dimensional (518 nm and 554 nm dual-wavelength) clustering plot option in the Bio-Rad QuantaSoft™ program.

### Intra-organismal selection assay

The strength of intra-organismal (within-host) selection on mutant mtDNA was measured longitudinally across isolated parent-progeny lineages, as previously described.^26^ Briefly, multiple L4-stage (late larval) heteroplasmic animals were picked at random under a dissecting microscope from stock populations carrying each heteroplasmy that had been crossed into the Bristol strain (wildtype) nuclear background. These larvae were transferred to fresh NGM plates seeded with live OP50 *E. coli* bacteria as a food source and incubated for 2 days at 20°C to allow adult maturation. The day-2 adults were individually segregated by transferring each onto a fresh food plate and incubated for 4 hours at 20°C to produce embryos that are age-synchronized to within a four-hour window. Each parent was then individually lysed. After 4 days of continued incubation at 20°C, the progeny had progressed from embryos to day-2 adults, reaching the same age at which their respective parents were lysed. Adult progeny were lysed at this point to obtain progeny that are age-matched to their parents, to control for age-dependent differences in mutant mtDNA levels. Progeny from each parent were lysed in pools of 3 to minimize the confounding effect of random drift. Each parent-progeny lineage was individually segregated from the rest, to ensure that mutant mtDNA frequency from each progeny lysate was being compared with that of its own respective parent, thereby minimizing the confounding effect of competition between lineages (organismal selection). Mutant mtDNA frequency of parents and progeny was determined for each heteroplasmy using ddPCR as described above, across multiple replicate parent-progeny lineages for statistical power.

### Experimental evolution (organismal selection)

Selection against mutant mtDNA that occurs strictly at the level of host fitness was measured using an organismal competition experiment similarly to the experiment previously described ^26^. Briefly, for each mutant mtDNA variant, heteroplasmic nematodes carrying mutant mtDNA in the Bristol nuclear background were combined with Bristol-strain nematodes on 10-cm NGM plates seeded with live OP50 *E. coli* bacteria as a food source. Approximately 500 nematodes were transferred to each plate. In addition to 4 replicate competition lines for each heteroplasmy, 4 non-competing control lines were established by transferring only heteroplasmic animals onto their own food plates, with no homoplasmic-wildtype animals to compete against. Every 3 days, the generation for each experimental line was reset; nematodes were washed off the plates using M9 buffer into a sterile 1.7 mL collection tube. Approximately 500 animals of mixed age from each line were transferred to a fresh food plate. Another 500 nematodes were lysed together in a single pooled lysate. This experiment was continued for 6 consecutive generations.

### Population genetic model

To treat the evolutionary dynamics of multilevel selection and drift in a theoretical population genetic framework, we constructed a model that couples a stochastic, frequency-dependent Wright-Fisher model for the evolution of mutant mtDNA frequencies within individuals together with a frequency-independent, deterministic model of organismal competition. The intra-organismal fitness function measures the fitness of mutant mtDNA relative to wildtype mtDNA within an individual. We model the intra-organismal fitness function as a sigmoid function of mutant frequency:

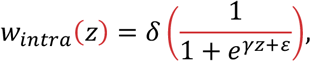

where parameters δ, γ, and ε control the overall scale of fitness variation, the degree of frequency-dependence, and the position of the inflection point, respectively, as a function of mutant mtDNA frequency. The organismal fitness function is modeled in terms of the fitness cost of carrying a selfish mitochondrial genome at frequency *z*:

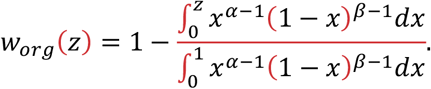

This is identical to the complementary cumulative distribution function of a beta distribution with parameters α and β, so that we necessarily have *w*_*org*_(0)=1 (wildtype individuals have fitness 1) and *w*_*org*_(1)=0 (individuals fixed for the selfish mutant are either sterile or nonviable). Importantly, when β=1 this expression reduces to *w*_*org*_(*z*)=1–*z*^α^, corresponding to a fitness defect that scales as *z* to the power α, or a linear fitness decline when both α=1 and β=1. This expression can thus encompass a variety of simple models for how mutant mtDNA frequency impacts organismal fitness, in addition to more complex threshold-like models. Note also that while the organismal fitness function depends on the mutant mtDNA frequency of the individual, in our model organismal selection is frequency-independent in that the fitness of an individual does not depend on the composition of the remainder of the population.

To combine this intra-organismal and organismal selection with stochastic inheritance arising from intra-organismal genetic drift, we construct an *N*+1 by *N*+1 matrix,

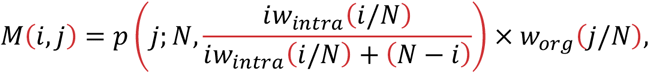

where the indices *i* and *j* run from 0 to *N, p*(*j*; *N, q*) is the binomial probability of a total of *j* successes out of *N* attempts with expected mutant mtDNA frequency *q* among the offspring of a parent with mutant mtDNA frequency *i*/*N* (which itself is given by *i w*_*intra*_(*i*/*N*) / (*i w*_*intra*_(*i*/*N*)+(*N*–*i*))). Thus *M*(*i, j*) gives the expected proportion of progeny with a mutant mtDNA frequency of *j*/*N* for a parent with a mutant mtDNA frequency *i*/*N*. Writing the probability distribution of heteroplasmic mutant frequencies at time *t* as a length *N* vector *f*_*t*_, the time evolution of this probability distribution is given by:

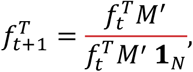

where *M*’ is the *N* by *N* principal submatrix of *M* obtained by omitting the zeroth row and column, and **1**_*N*_ is the length *N* vector of all 1s. By the Perron-Frobenius theorem,^63^ as *t* goes to infinity, *f*_*t*_ converges to a stable distribution *f*\* given by the dominant left eigenvector of *M*’. The corresponding eigenvalue gives the expected number of heteroplasmic offspring per heteroplasmic parent under the stable distribution *f*\*. The expected number of wildtype offspring per heteroplasmic parent (spontaneous reversion from heteroplasmy to homoplasmic-wildtype) under the stable distribution *f*\* is given by:

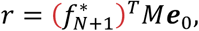

where *f*\*_*N*+1_ is the vector of length *N*+1 whose zeroth entry is 0 and whose remaining entries are given by *f*\*, and ***e***_0_ is the length *N*+1 vector whose zeroth entry is 1 and whose remaining entries are 0.

### Maximum likelihood inference

We estimate the model parameters for each mutant mtDNA genotype using a joint maximum likelihood approach (that is, a combined fit to the data from the intra-organismal selection experiment, organismal selection experiment, and the sampled stable mutant frequency distribution) with parameter uncertainty assessed via a parametric bootstrap approach. Model fitting was conducted using a custom Python script (code available at https://github.com/bgitschlag/MiSelf).

For intra-organismal selection experiment, we model the mutant frequency within the progeny, *z*_*obs,t*+1_, as a function of the mutant frequency *z*_*t*_ of the parent as:

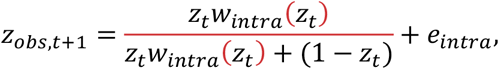

where *e*_*intra*_ is a normally distributed random variable with mean 0 and variance given by the free parameter σ^2^_*intra*_. For the organismal selection experiment, the population-wide fraction of heteroplasmic individuals, *ϕ*, changes across generations due to both the organismal fitness cost and spontaneous loss of the mutant genome, *r*. At each generation, *ϕ* experiences a growth rate, relative to the wildtype, of *κ*=(*f\**)^*T*^*M*’**1**_*N*_, and a rate of mutant loss *r*, which collectively determine the fraction of heteroplasmic individuals in each subsequent generation:

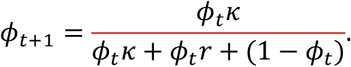

Because in the special case where mutant frequency declines solely due to organismal selection (that is, *r*=0), ln(*ϕ*/(1–*ϕ*)) decreases linearly in time, we model *ϕ* on a log-odds scale:

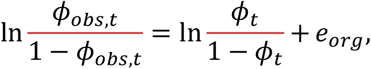

where *ϕ*_*obs,t*_ is the empirically observed *ϕ* at time *t, e*_*org*_ is a normally distributed random variable with mean 0 and variance given by the free parameter σ^2^_*org*_. In addition, the initial frequency *ϕ*_*0*_ is also inferred as a free parameter. Finally, to determine the likelihood of draws from *f*\*, we construct a continuous analog of the discrete distribution *f*\* by applying a Gaussian smoother and calculating the likelihood under the resulting probability density function. Specifically, we convolve the discrete distribution described by *f*\* with a Gaussian random variable with mean 0 and standard deviation 1/*N*.

In summary, our statistical model has free parameters, *N*, δ, γ, and ε (controlling intra-organismal selection and drift), σ^2^_*intra*_ (controlling the noise variance of the intra-organismal selection experiment), α and β (controlling the form of organismal selection), and σ^2^_*org*_ (controlling the noise variance in the organismal selection experiment). Because all of these parameters are continuous except for *N*, for each possible *N* from 10 to 100 we numerically maximized the likelihood with respect to all other parameters. We then selected the value of *N* that maximized the likelihood.

To estimate our confidence in these parameter estimates, we performed parametric bootstrapping by simulating 100 data sets per genotype, using the sample sizes, maximum-likelihood population-genetic parameters, and error estimates for the corresponding empirical data set. Specifically, to simulate the intra-organismal selection experiment, parent mutant frequencies *z*_*t*_ were randomly drawn from the Gaussian smoothed version of *f*\* and then the corresponding progeny frequencies *z*_*t*+1_ were randomly drawn according to our inferred Wright-Fisher process. To simulate the organismal selection experiment, the expected heteroplasmic fraction *ϕ*_*t*_ at each generational time *t* was calculated using the maximum-likelihood estimate of *ϕ*_0_ combined with the *r* and *κ* values determined from our maximum-likelihood estimates. We then converted these expected fractions *ϕ*_*t*_ onto a log odds scale, added Gaussian noise with variance σ^2^_*org*_, and finally converted them back to a frequency scale. The samples from the mutant frequency distribution were simulated by drawing from the Gaussian smoothed version of *f*\* with sample sizes equaling that of the corresponding empirical data set. For each simulated data set, model parameters were re-estimated using the same maximum-likelihood approach described above. However, for some bootstrap simulations the maximum likelihood estimate of *N* was realized at or near the upper boundary of our search space. In such cases we thus examined larger values of *N*, starting at *N*=125 and increasing by powers of 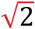 (125, 177, 250, 354, 500, etc.). We continued in this manner until the maximum-likelihood estimate of *N* reached a peak, defined as a maximum-likelihood *N* that is no longer among the 2 consecutive largest values of *N* examined so far, or until reaching a maximum-likelihood *N*=1,000, at which point we concluded that the magnitude of drift approximates that of the high-*N* limit.

## Supporting information

mpt2_intra_organismal_selection_source_source_data

mpt2_organismal_selection_source_data

mpt4_intra_organismal_selection_source_data

mpt4_organismal_selection_source_data

mptDf2_intra_organismal_selection_source_data

mptDf2_oganismal_selection_source_data

mptDf3_intra_organismal_selection_source_data

mptDf3_organismal_selection_source_data

mutant_frequency_samples_source_data

uaDf5_intra_organismal_selection_source_data

uaDf5_organismal_selection_source_data

## DATA AVAILABILITY

Data from experiments measuring selection on *uaDf5* were previously published.^26^ These and other experimental data encompassing the remaining four genotypes are included as Source Data files in this study.

## CODE AVAILABILITY

All code is publicly available at https://github.com/bgitschlag/MiSelf

## ACKNOWLEDGMENTS

The *uaDf5* strain was kindly provided by the Caenorhabditis Genetics Center (CGC). We thank O. Thompson and R. Waterston at the University of Washington for identifying the *mptDf2, mpt4, mpt2*, and *mptDf3* genotypes from the Million Mutation Project worm collection. We thank Patrick Abbot and Ann T. Tate of Vanderbilt University for valuable conceptual insights and guidance with experimental design. This work was supported by the NIH research project grants R01 GM123260 and R35 GM145378 (M.R.P.), Discovery Award (PR170792) from the Department of Defense’s Congressionally Directed Medical Research Program (M.R.P.), the Vanderbilt University Medical Center Diabetes Research and Training Center Pilot & Feasibility Grant (M.R.P.), the NIH-sponsored Cellular, Molecular and Biochemical and Molecular Sciences Training Program (5T32GM008554-18, B.L.G.), the Ruth L. Kirschstein National Research Service Award Individual Predoctoral Fellowship (1F31GM125344, B.L.G.), R35 GM133613 (D.M.M.), an Alfred P. Sloan Research Fellowship and additional funding from the Simons Center for Quantitative Biology and Cold Spring Harbor Laboratory (D.M.M.).

## AUTHOR CONTRIBUTIONS

B.L.G., D.M.M., and M.R.P. designed the study. B.L.G., C.V.P., and J.P.H. performed experiments to collect the data. B.L.G. and D.M.M. performed mathematical modeling. B.L.G., D.M.M., and M.R.P. analyzed the data. B.L.G. prepared the first manuscript draft. All authors revised and edited the manuscript.

## ADDITIONAL INFORMATION

Correspondence should be addressed to Bryan L. Gitschlag or Maulik R. Patel.

