## Supplementary_information for "Multiple distinct evolutionary mechanisms govern the dynamics of selfish mitochondrial genomes"

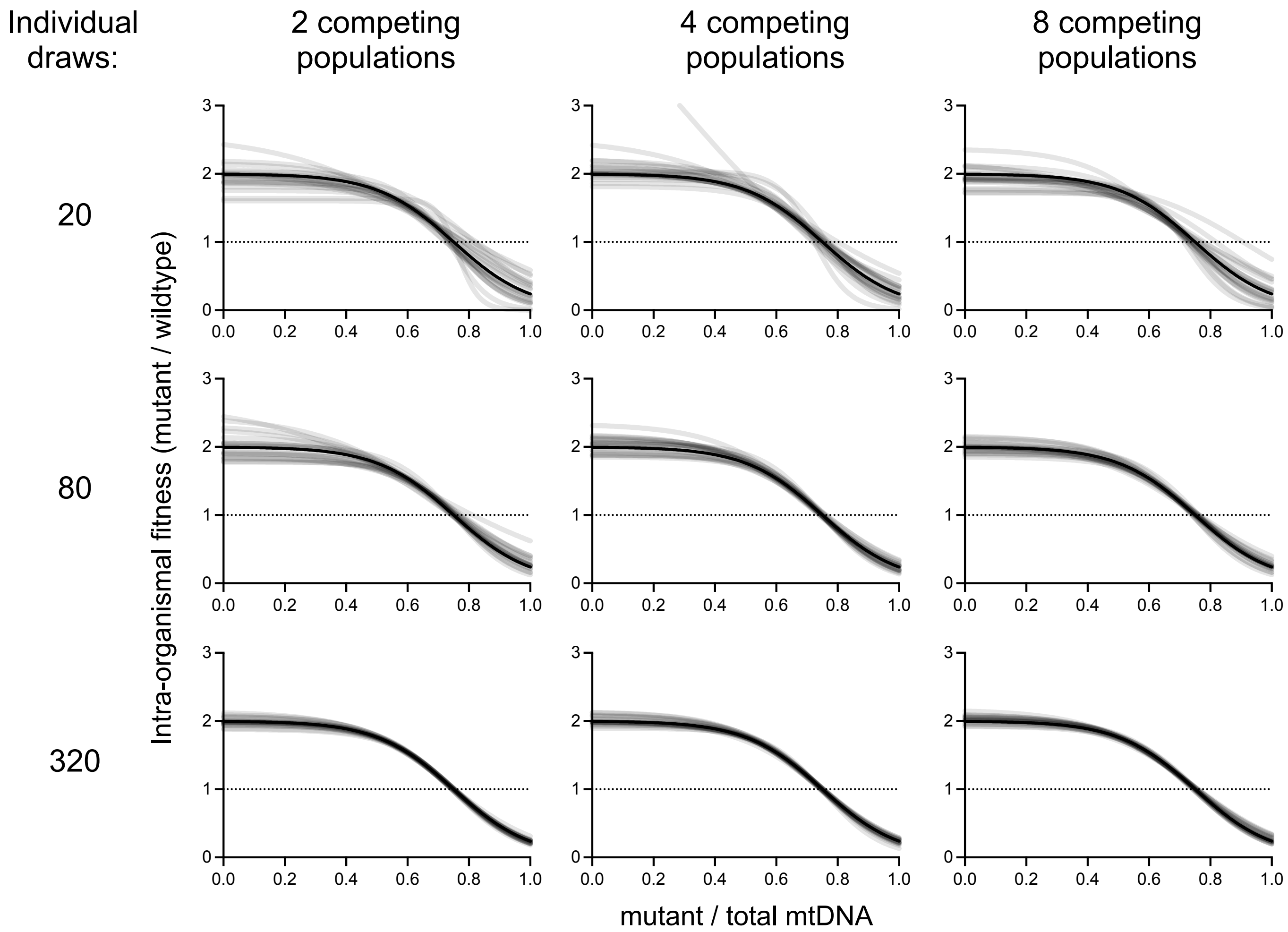


**Supplementary Figure 1: Intra-organismal fitness functions for simulated data of different sample sizes.**

Intra-organismal fitness functions obtained by simulating data of different sample sizes (n=20 simulations per plot) from a single set of unique input model parameters. The true fitness function for the model parameters is shown as a solid black line on each plot. Model parameters were re-estimated by joint maximum likelihood for each simulated data set, resulting in a unique combination of a new intra-organismal fitness function (gray lines), organismal fitness function (Supplementary Fig. 2), and most evolutionarily stable mutant frequency distribution (Supplementary Fig. 3). The data points drawn from the mutant frequency distribution vary by row (increasing from top to bottom). These are used as samples of the mutant frequency distribution and as the parents for modeling intra-organismal selection. The number of replicate populations, simulated for the purposes of modeling organismal selection, vary by column (increasing from left to right).

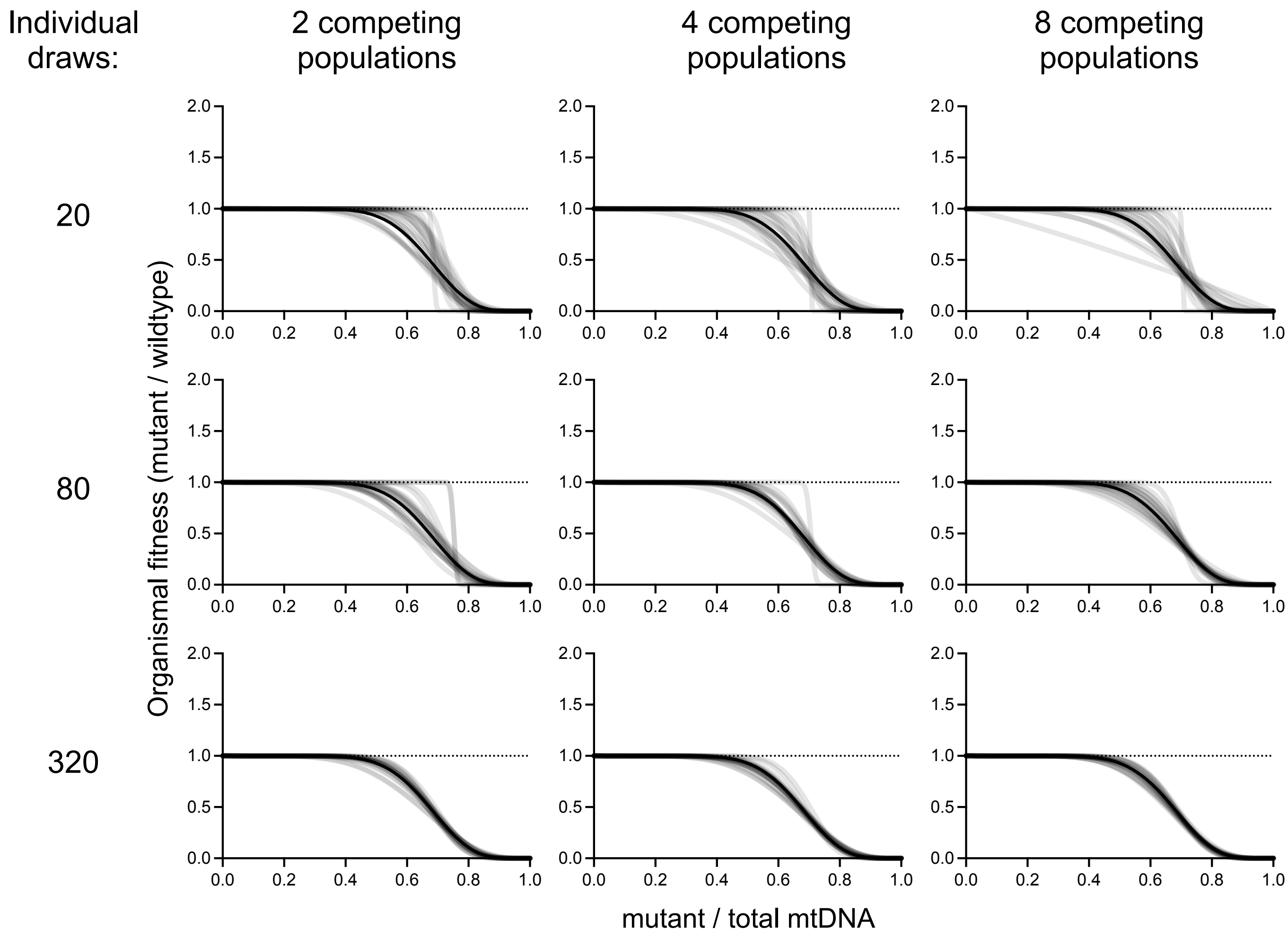


**Supplementary Figure 2: Organismal fitness functions for simulated data of different sample sizes.**

Organismal fitness functions obtained by simulating data of different sample sizes (n=20 simulations per plot) from the same set of unique input model parameters as used in Supplementary Fig. 1. The true fitness function for the model parameters is shown as a solid black line on each plot. Model parameters were re-estimated by joint maximum likelihood for each simulated data set, resulting in a unique combination of a new intra-organismal fitness function (Supplementary Fig. 1), organismal fitness function (gray lines), and most evolutionarily stable mutant frequency distribution (Supplementary Fig. 3). The data points drawn from the mutant frequency distribution vary by row (increasing from top to bottom). These are used as samples of the mutant frequency distribution and as the parents for modeling intra-organismal selection. The number of replicate populations, simulated for the purposes of modeling organismal selection, vary by column (increasing from left to right).

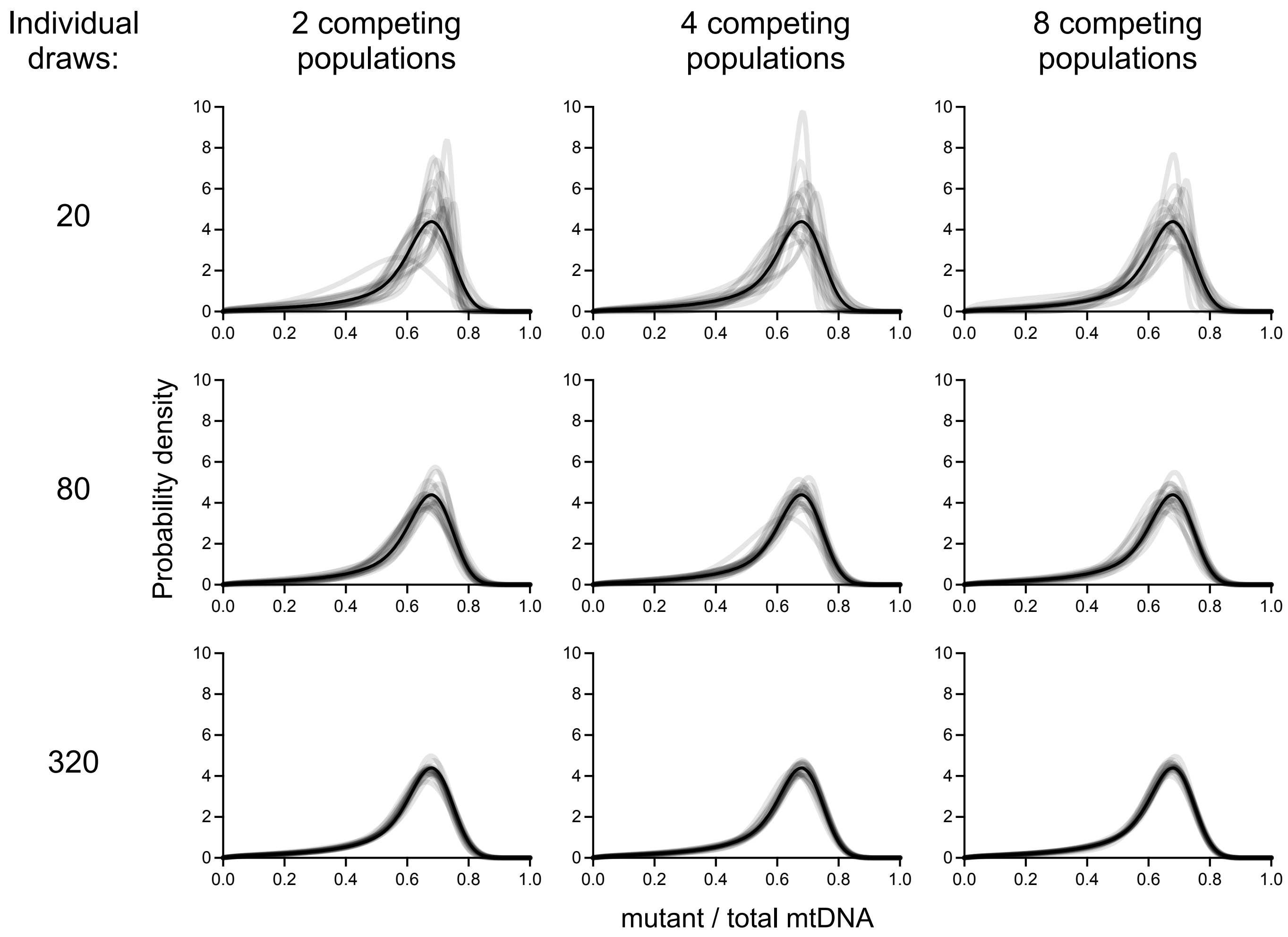


**Supplementary Figure 3: Stationary distributions for simulated data of different sample sizes.**

Stationary distributions obtained by simulating data of different sample sizes (n=20 simulations per plot) from the same set of unique input model parameters as used in Supplementary Fig. 4 and 5. The true fitness function for the model parameters is shown as a solid black line on each plot. Model parameters were re-estimated by joint maximum likelihood for each simulated data set, resulting in a unique combination of a new intra-organismal fitness function (Supplementary Fig. 1), organismal fitness function (Supplementary Fig. 2), and the most evolutionarily stable mutant frequency distribution (shown here in gray). The data points drawn from the mutant frequency distribution vary by row (increasing from top to bottom). These are used as samples of the mutant frequency distribution and as the parents for modeling intra-organismal selection. The number of replicate populations, simulated for the purposes of modeling organismal selection, vary by column (increasing from left to right).

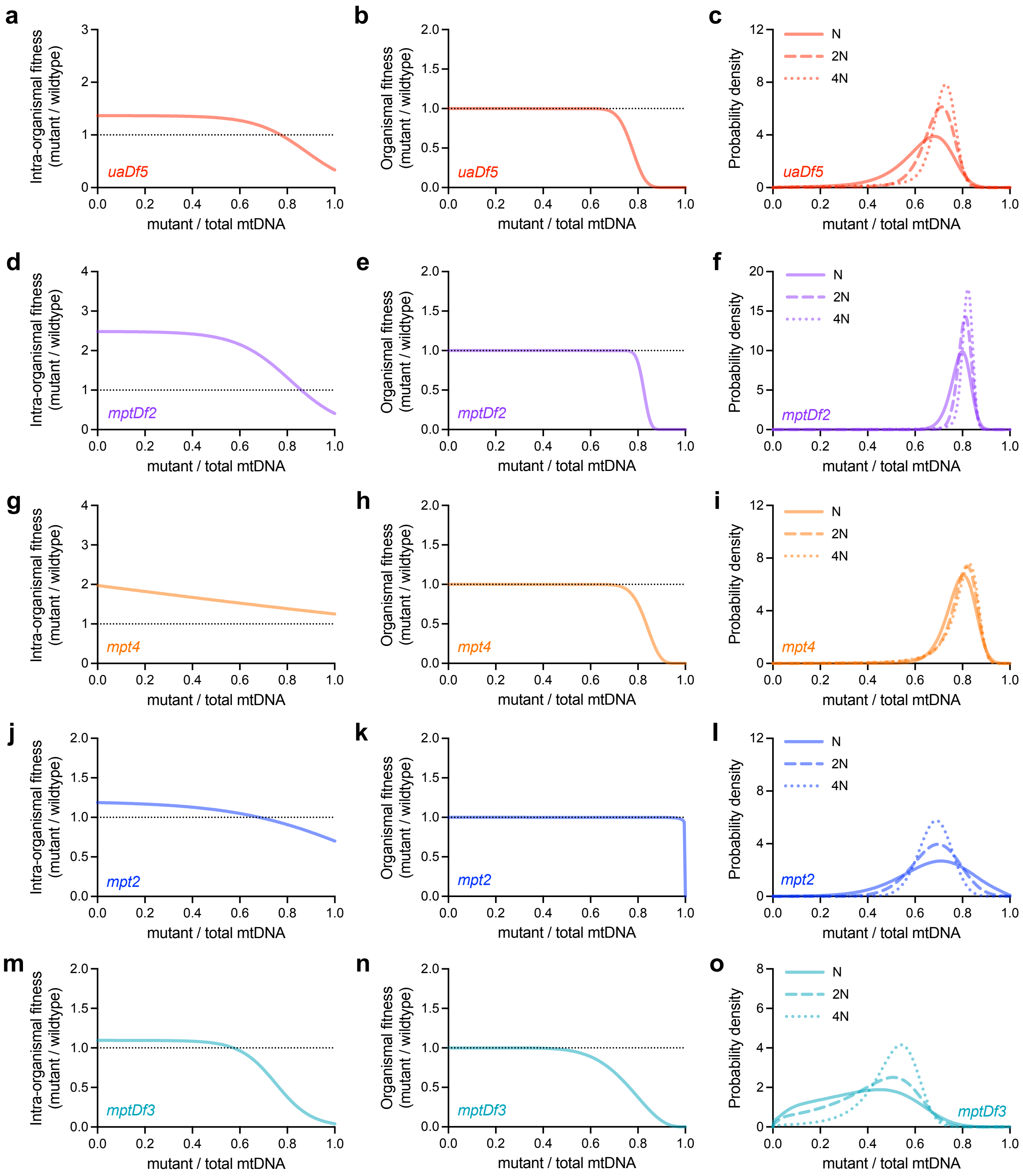


**Supplementary Figure 4: Effect of variation in magnitude of drift on the mutant mtDNA frequency distribution under the maximum-likelihood selection parameters.**

Maximum likelihood estimates of the intra-organismal (left column) and organismal (center column) fitness effects, each as a function of mutant frequency, similar to Figure 5 but excluding bootstraps, and the most evolutionarily stable mutant frequency distribution (right column) under maximum-likelihood selection parameters and the drift parameter (N), in addition to two- and four-fold changes in magnitude of drift (2N and 4N, respectively).

| Genotype | Generations until probability of invasion >0.5 | | | |
|---|---|---|---|---|
| | $N_e$=10 | $N_e$=100 | $N_e$=1,000 | $N_e$=10,000 |
| *uaDf5* | 287 | 29 | 3 | <1 |
| | 121–$3.4*10^4$ | 12–3,418 | 1–342 | <1–34 |
| *mptDf2* | $6.2*10^{15}$ | $6.2*10^{14}$ | $6.2*10^{13}$ | $6.2*10^{12}$ |
| | 3,296–$1.9*10^{24}$ | 330–$1.9*10^{23}$ | 33–$1.9*10^{22}$ | 3–$1.9*10^{21}$ |
| *mpt4* | $2.0*10^7$ | $2.0*10^6$ | $2.0*10^5$ | $2.0*10^4$ |
| | 2,791–$9.4*10^{18}$ | 279–$9.4*10^{17}$ | 28–$9.4*10^{16}$ | 3–$9.4*10^{15}$ |
| *mpt2* | $1.1*10^5$ | $1.1*10^4$ | 1,094 | 109 |
| | 531–$7.9*10^{10}$ | 53–$7.9*10^9$ | 5–$7.9*10^8$ | 1–$7.9*10^7$ |
| *mptDf3* | 24 | 2 | <1 | <1 |
| | 7–3,929 | 1–393 | <1–39 | <1–4 |

**Supplementary Table 6: Expected persistence times of the heteroplasmic state.**

Persistence or robustness of the heteroplasmic state, defined by the expected number of generations until the invasion by a de novo homoplasmic lineage (introduced by the loss of the selfish genome through intra- organismal drift) becomes more probable than no invasion. Since the loss of the mutant genome is beneficial to host fitness, the probability of invasion is treated as the fixation probability for a beneficial genotype, described in ref. 48. Briefly, the per-generation probability of invasion is $1\text{-}e^{-2spNe}$, where the selection coefficient s is equal to w-1, where w is the relative fitness of homoplasmic organisms, $1/w_{org}$. Here, p is the per-generation probability of de novo introduction of a homoplasmic lineage (equal to the probability of mutant loss, provided in Table 1 column titled probability of loss), and Ne is the effective organismal population size. Since the per-generation probability of no invasion is equal to $e^{-2spNe}$, the persistence time as defined is thus equal to the minimum value of generations g such that $(e^{-2spNe})g \leq 0.5$. Each entry: persistence time given maximum-likelihood model parameters (first line), in addition to 95% bootstrap confidence interval (second line).

| Genotype | Average *z* | *z** |
|---|---|---|
| *uaDf5* | 0.63 | 0.77 |
| | 0.59–0.65 | 0.72–>1 |
| *mptDf2* | 0.79 | 0.86 |
| | 0.77–0.80 | 0.83–0.93 |
| *mpt4* | 0.79 | >1 |
| | 0.77–0.81 | 0.91–>1 |
| *mpt2* | 0.67 | 0.68 |
| | 0.61–0.73 | 0.62–0.84 |
| *mptDf3* | 0.38 | 0.57 |
| | 0.30–0.45 | 0–0.80 |

**Supplementary Table 7: Comparison of average heteroplasmic mutant mtDNA frequency and heteroplasmic frequency predicted by intra-organismal balancing selection (z*).**

Each entry: maximum likelihood estimate (first line), 95% bootstrap confidence interval (second line).
